# VIP+ amacrine cells synchronize neural activity in the retina

**DOI:** 10.64898/2026.02.06.704482

**Authors:** Déborah Varro, Samuele Virgili, Simone Ebert, Gabriel Mahuas, Ulisse Ferrari, Olivier Marre

**Affiliations:** Institut de la Vision, Sorbonne Université, INSERM, CNRS, Paris, France

## Abstract

In early sensory circuits, inhibitory interneurons are best known for mediating lateral inhibition, gain control, and feature selectivity, but their roles at the population level are less understood. Using two-photon digital holography, we found that in the retina, vasoactive intestinal peptide–expressing (VIP^+^) amacrine cells not only inhibit some retinal ganglion cell types via GABAergic synapses but also excite specific other types through gap junctions. Selective knockout of these junctions abolished synchronized firing among these ganglion cells, showing that VIP^+^ amacrine cells coordinate their activity. Thus, VIP+ interneurons have a dual, cell–type–specific role: synchronizing some ganglion cells electrically while inhibiting others chemically, thereby differentially shaping retinal output.

## Introduction

Inhibitory interneurons form a diverse population in sensory areas and substantially reshape the responses of excitatory cells (1). In the early visual system, a major role of these interneurons is to perform lateral inhibition: they receive inputs from local excitatory cells and inhibit the more distant excitatory cells of the same type. This implements surround suppression in various areas (2,3). Inhibitory interneurons are also involved in gain control, normalizing the output based on the past history of the input (4). Overall, they allow sensory responses to integrate the stimulation context. Finally, inhibitory interneurons are also involved in feature selectivity, for example, in the visual cortex, where they contribute to orientation tuning (5). While these roles have been extensively explored at the single-cell level, it remains unclear whether they have additional functions that shape population activity.

In the retina, the visual signal sent to the brain by retinal ganglion cells (RGCs) depends on inhibitory interneurons, particularly amacrine cells (ACs). Blocking the transmission of ACs significantly changes the response of GCs and thus the visual information sent to the brain, demonstrating their importance in visual signal processing (6,7).

ACs exhibit a great diversity, with 60 different types identified in mice, based on morphological and genetic classifications (8–11). Both their morphological and physiological diversity allow them to tune and modify the visual signal sent to the GCs in various ways (12). Despite their importance in the retina, the impact of many types of ACs on GCs remains unclear, especially at the population level.

A major class of inhibitory interneurons is the VIP+ interneurons, which have been extensively studied in the cortex. ACs that express vasoactive intestinal polypeptide (VIP+) can be targeted by using a Cre-expressing transgenic mouse line under the control of the VIP promoter (13). There are three types of VIP+ ACs in the mouse retina: VIP-1, VIP-2, and VIP-3, which differ in their morphology (14–17) and light sensitivity (9,10). VIP-2 and VIP-3 are small and narrow medium-field cells in the Inner Nuclear Layer (INL) and Ganglion Cell Layer (GCL), while VIP-1 ACs are wide-field, bistratified cells in the INL. VIP-1 ACs have long ramifications and project into both the OFF and ON layers of the Inner Plexiform Layer (IPL), allowing them to contact both ON and OFF GCs (15–17). Physiologically, all three VIP+ AC types depolarize in response to ON stimuli and hyperpolarize to OFF stimuli, showing selectivity based on stimulus size (15). VIP+ cells are especially interesting because anatomical studies have shown that they form both inhibitory GABAergic synapses (15,16) and excitatory electrical synapses through gap junctions (15). VIP+ ACs populations can inhibit some types of GCs via GABAergic signaling when all the cells of the population are simultaneously stimulated with full-field optogenetic stimulation (15). However, it is unclear whether a single VIP+ cell can inhibit GC activity. In addition to their inhibitory synapses, VIP-1 cells form excitatory electrical synapses via gap junctions with other VIP-1 cells and with other targets whose identity is unclear (15,17). The functional effects of these excitatory electrical synapses on GCs remain unexplored.

To study how VIP+ ACs influence GC activity and their functional role in retinal visual signal processing, we combined 2-photon holographic optogenetic stimulation with electrophysiological recordings. We used a transgenic mouse line in which VIP+ cells express Cre recombinase (13–16) and were made light-sensitive by expressing a depolarizing optogenetic protein under the control of Cre (Fig. 1A,1B). We stimulated VIP+ cells individually using two-photon digital holography (18,19). At the same time, we recorded the impact of single VIP+ cell stimulations on GCs with a multi-electrode array (MEA) (20,21). This method allowed us to determine how a single VIP+ AC can modulate the activity of different GC types.

**Fig 1.**
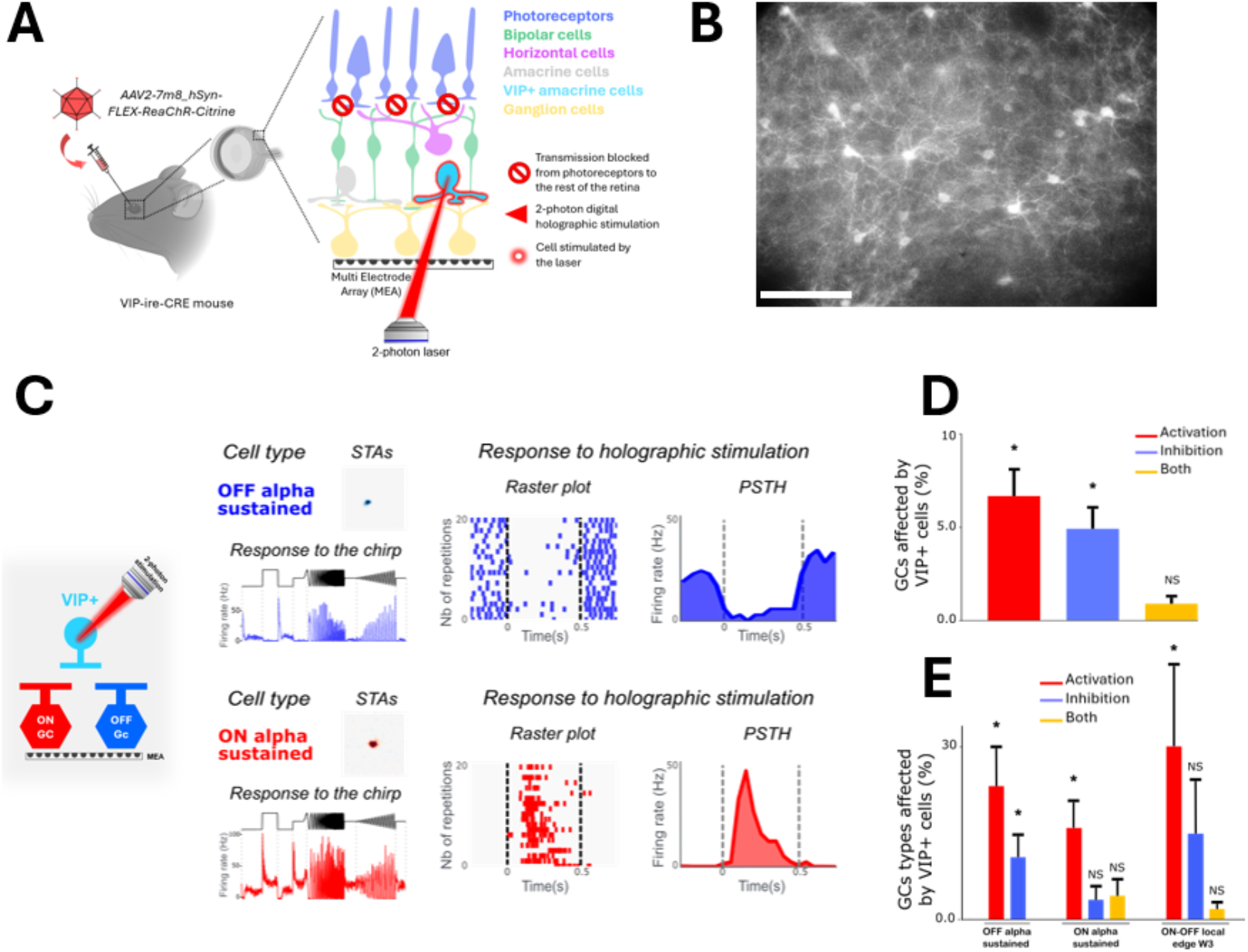
VIP+ cells are capable of both inhibiting and activating ganglion cells. (A). Schematic representation of recording GCs with a multi-electrode array and simultaneous 2-photon holographic stimulation of VIP+ cells expressing ReaChR-Citrine after LAP4+ACET application (5 and 1 µM). (B) Epifluorescent picture of a retina piece expressing VIP+ ACs coupled with a fluorescent protein, showing their soma and ramifications. The scale bar is 100 microns. (C). Responses (raster plots and PSTHs) of two GCs to the holographic stimulation of one individual VIP+ cell. The GCs are from different types (ON & OFF), as indicated by the polarity of their STAs and their responses to the chirp stimulus. The holographic stimulation duration is indicated by dashed lines (500 ms, Pmax = 0.5 mW for 1040 nm). (D). Percentages of GCs affected by VIP+ cell stimulation. (E). Percentages of GCs affected by VIP+ ACs stimulation by type. For (D) and (E), the values reported in the bar plots are mean ± S.D.M. across experiments. Values are considered significant (*) if the mean is at least two *σ*s larger than zero.

We found that VIP+ ACs could both inhibit specific OFF types through GABAergic inhibition but also excite other GC types, including ON alpha and W3-like, through their gap junctions. By knocking out these gap junctions specifically, we found that they allow VIP+ ACs to synchronize these specific types of GCs. Our results thus uncover a dual role of VIP+ amacrine cells, which extends beyond inhibition, by synchronizing specific types of GCs at the population level.

## Results

### VIP+ amacrine cells can both inhibit and activate ganglion cells

To understand how VIP+ ACs modulate the activity of different types of RGCs, we conducted experiments combining optogenetic stimulation of VIP+ ACs with electrophysiological recordings of GCs. We used VIP-ires-Cre mice (13–16,22) in which the VIP+ ACs were made sensitive to light by expressing the optogenetic protein ReaChR coupled with a green fluorescent protein by AAV injection. We recorded GC activity in the retinas using multi-electrode arrays (MEAs) (20,21). We first measured the receptive fields (RFs) of GCs, their types, and orientation/direction selectivity by recording their responses to a checkerboard stimulus, a chirp, and drifting gratings, following the method of Baden et al (2016) (23). To perform optogenetic stimulation of VIP+ ACs without stimulating the retinal circuit through photoreceptors, we blocked their transmission to the rest of the retina using LAP-4 (5 µM) and ACET (1 µM). To measure the effect of each VIP+ cell separately on the GCs recorded, we used 2-photon digital holography to stimulate with a laser beam (1040λm, 500ms, 20reps) each VIP+ AC individually (18) (see schematic representation of the method in **Fig.1A**).

We found that VIP+ ACs can both inhibit and activate different types of ON and OFF GCs. Specifically, OFF alpha-sustained GCs were significantly suppressed by the optogenetic stimulation of VIP+ ACs (11% ± 3.8 of all recorded OFF alpha-sustained GCs showed reduced activity) (**Fig. 1C,1E**). This finding aligns with previous research using full-field optogenetic stimulation (15). Conversely, we also observed that VIP+ ACs can activate specific types of GCs. The GCs significantly activated by VIP+ ACs included OFF alpha-sustained, ON alpha-sustained, and ON-OFF local-edge W3 GCs, with 23% ± 6.5, 16% ± 4.5, and 35% ± 17 of recorded cells affected respectively (**Fig. 1C,1E**). Among all recorded GCs (n = 2,959 from 18 retinas), 4.9% were significantly inhibited, and 6.7% were significantly activated by the holographic stimulation of VIP+ ACs (**Fig. 1D**). These findings demonstrate that VIP+ ACs have distinct and type-specific inhibitory or excitatory effects on both ON and OFF GCs.

These results altogether showed that one VIP+ AC is enough to modulate alone the activity of GCs (**Fig.1C**). We also observed that a single VIP+ AC can modulate the activity of several GCs, and that several VIP+ ACs can contact the same GC (**Supp.Fig. 1**). Moreover, we found that some VIP+ ACs could simultaneously activate some GCs while inhibiting other GCs (ex, **Fig. 1C**). This suggests that the VIP+ ACs that mediate the activation and inhibition are not divided in two distinct pools, but rather that the two modes of activation coexist in a single VIP cell.

### VIP+ amacrine cells inhibit ganglion cells via GABA signalization and excite other ganglion cells via gap junctions

We then aimed to understand the mechanisms that allow VIP+ ACs to excite some types of GCs and inhibit others. VIP+ cells are GABAergic interneurons that not only co-stain for GABA but also make functional GABAergic inhibitory synapses with GCs (24,13–16,25,22). To determine if both the inhibition and the activation by VIP+ ACs were governed by GABA signalization, we compared the GC responses during the optogenetic stimulation of VIP+ ACs, before and after blocking GABA transmission. We performed separate experiments using different pharmacological blockers to block GABA-A receptors (SR95531, 20 µM) and GABA-C receptors (TPMPA, 10 µM) individually. The activity of GCs was not significantly affected after applying the GABA-C receptor blocker (p > 0.5, n=2 retinas) (**Fig.3A and 3B**), indicating that VIP+ ACs’ modulation of GCs does not depend on GABA-C receptors. Conversely, the number of GCs inhibited by the stimulation of VIP+ ACs decreased substantially after blocking GABA-A receptors (more than 70% of GCs lost their inhibition). In contrast, the number of GCs activated by VIP+ cells was significantly less affected by the drug application (75% of them remained activated) (n = 6 retinas; p < 0.001) (**Fig.2A and 2B**). The observed 25% loss may be attributed to the experiment’s duration, which may have made some cells unresponsive.

Our results showed that VIP+ ACs inhibit GCs through GABAergic transmission and GABA-A receptors. However, the activation effect of VIP+ ACs on GCs is not solely mediated by GABAergic signaling. This result rules out the hypothesis that the activation might be due to disinhibition, meaning VIP+ ACs inhibit an AC that would then disinhibit GCs. If that were the case, blocking GABA transmission would also block this disinhibition. Therefore, the activation effect of VIP+ cells on GCs depends on an alternative mode of transmission.

VIP-1 ACs have been shown to form gap junctions between neighboring VIP+ ACs of the same type (homologous couplings) and with other types of cells (heterologous couplings) (15). We thus tested whether gap junctions could mediate the observed excitatory responses of GCs. We compared GC responses to optogenetic stimulation of VIP+ ACs before and after blocking gap junctions (18BGa, 25 µM). Our results confirm that blocking gap junctions significantly reduced the responses of GCs activated by VIP+ ACs stimulation (only 4.5% ± 3.9% remained activated), compared to the responses of GCs that were inhibited (n = 4 retinas, p < 0.05) (**Fig.2C and 2D**). This demonstrates that the activation effect of VIP+ ACs is mediated through gap junctions.

To further determine the exact pathway by which VIP+ ACs can activate some types of GCs, we tested whether their excitatory responses were dependent on glutamatergic transmission. We performed the same experiments as before while blocking glutamatergic receptors. We used pharmacological blockers of AMPA, kainate, and NMDA receptors (CNQX, 20 µM; CPP, 10 µM), a combination that blocks most of the glutamatergic transmission in the retina. GC responses evoked by the stimulation of VIP+ ACs were not significantly affected after the addition of this pharmacological cocktail (n = 1 retina; p > 0.2) (**Fig.3C and 3D**). Taken together with the previous findings, this shows that the GCs are activated by a pathway that does not rely on glutamatergic transmission, but on gap junctions. This suggests that GCs activated by VIP+ ACs are directly connected with them via gap junctions, rather than through bipolar cells.

The observed inhibitory effect of VIP+ ACs on GCs could be mediated by direct GABAergic synaptic inhibition or, VIP+ ACs might inhibit bipolar cells, thereby reducing their excitatory input to GCs. To distinguish these two hypotheses, we examined the effect of blocking glutamatergic transmission on inhibitory GC responses. Our results show that blocking glutamatergic transmission did not significantly alter the inhibitory responses of GCs evoked by stimulating VIP+ ACs (p > 0.2) (**Fig.3C and 3D**). This aligns with previous findings showing that VIP+ ACs form direct GABAergic synapses with GCs (15,26–28) and rules out the hypothesis that VIP+ ACs inhibit bipolar cells to suppress their glutamatergic excitation of GCs.

Because VIP+ ACs express the VIP neuromodulator, we tested whether VIP modulates GC responses. We used the same protocol as before while blocking VIP receptors in the retina (VPAC1 and VPAC2) with a pharmacological blocker ([Ac-Tyr1, D-Phe2]GRF, 300 nM)(29,30). The activation and inhibition responses of GCs evoked by VIP+ AC stimulation were not significantly affected after blocking the VIP receptors (n=2 retinas, p > 0.2) (**Fig.3E and 3F**). The VIP neuromodulator does not appear to change the way VIP+ cells modulate GCs in the retina, although it might have effects on the rest of the retinal network that could not be visible in our recordings (31,32).

Taken together, our findings demonstrate that VIP+ ACs can modulate different types of GCs in two opposing ways and through two distinct pathways: VIP+ ACs activate specific types of GCs via gap junctions, while inhibiting other GCs through GABAergic synapses.

### VIP+ amacrine cells synchronize specific types of ganglion cells

Gap junctions can synchronize the firing of neurons. Since GCs connected to the same AC through gap junctions tend to fire in synchrony (33,34), we hypothesized that VIP+ ACs might synchronize the activity of the GCs they connect via gap junctions. This hypothesis is further supported by the timescale of the noise correlation among cells of these types, which is compatible with an indirect connectivity scheme going through an amacrine cell (35) (**Fig. 4C**).

To confirm that VIP+ ACs play a role in synchronizing some types of GCs, we specifically suppressed the gap junctions in VIP+ ACs by crossing VIP-ires-Cre mice with Cx36-fl/fl mice. We obtained an offspring (referred to as VIP:KO:Cx36) in which connexin 36 is knocked out specifically in VIP+ ACs. We then compared the noise correlation (NC) of same-type GCs between the control mouse’s retina and the VIP:KO:Cx36 retina. We presented a checkerboard stimulus (over 60 repetitions) to activate GCs and measured the NC and noise covariance in both conditions for each pair of same-type GCs (see schematic in **Fig. 4A, 4B**). Our results showed that NC was significantly reduced in the KO condition (n=3 retinas) compared to the control (n=5 retinas) for ON alpha-sustained (p<5e-23) (**Fig. 4C, 4D, and 4F**) and ON-OFF local-edge-W3 GCs (p<5e-10) (**Figs. 4C, 4E, and 4F**). The number of pairs with significant NC for these two GC types also strongly decreased. Gap junctions between VIP+ ACs and ON alpha-sustained, and W3 GCs are thus essential for maintaining high NC and, consequently, synchrony among these cell types. These GC types are also those specifically excited by optogenetic stimulation of VIP+ ACs (**Fig. 1E**).

Could it be that all gap junctions were knocked out in our mouse model and not just the ones in VIP+ ACs? To test this, we measured the NC between OFF alpha-sustained GCs. These cells are known to be coupled through direct gap junctions, not going through ACs (36). As a result, the neighboring cells of this type have a cross-correlogram that shows a double peak around zero, which is characteristic of directly coupled cells (37,38). We confirmed these findings in our own data (**Fig. 4H**). OFF alpha-sustained GC pairs do not show any significant change in their NC between the WT and KO conditions (p > 0.1) (**Fig. 4I**). This shows that our mouse model, VIP:KO:Cx36, does not disrupt all gap junctions in the retina, as direct gap junctions between OFF alpha-sustained GCs are preserved.

To further confirm that gap junctions mediated the observed NCs, we applied a gap junction blocker (18βGa, 25 µM). This substantially and significantly reduced NCs, both in the types that are directly coupled without VIP, and those that were coupled through the VIP ACs (**Fig. 4G**). Taken together, our findings indicate that VIP+ ACs synchronize some types of RGCs, the ON alpha sustained and W3, through their gap junctions.

## Discussion

Amacrine cells in the retina are crucial for visual processing; however, the functional role and impact of most of them on GCs remain largely unexplored. A classical assumption is that they are involved in surround suppression and gain control. VIP+ ACs are known to be GABAergic inhibitory interneurons (15,22). A previous anatomical study showed that one type of VIP+ cells, the bistratified VIP-1 ACs, form gap junctions with other VIP-1 ACs and some unidentified cells (14). However, the functional effect of these gap junctions on the GC population was unclear, and their role in retinal visual processing was unknown. In this study, we showed that VIP+ ACs had a dual impact on GCs. They inhibit OFF alpha GCs but activate ON alpha and W3 GCs through gap junctions. These gap junctions create a connectivity in which ON alpha and W3 cells are coupled to their neighbor cells of the same type via VIP+ ACs. This mechanism is responsible for the NCs observed between these types. VIP+ ACs synchronize specific cell types through gap junctions, while inhibiting others with their GABAergic synapses.

This multifunctional connectivity can be reminiscent of AII ACs, which connect to ON bipolar cells via gap junctions and to OFF bipolar cells via glycinergic synapses. The difference here is that VIP+ ACs likely connect directly to ON GCs through gap junctions, not through bipolar cells. This enables VIP ACs to play a key role in the emergence of NCs between specific GC types.

Our results are consistent with a previous report showing that VIP ACs inhibit specific types of GCs (15). However, the activation of specific types through gap junctions has not been previously reported. Previous studies suggested this “connectivity triangle”, where a GC is connected through a gap junction to an AC, which is itself connected to another GC of the same type (39,40). This was supported by anatomical evidence (41); however, direct functional and causal evidence were lacking, and the AC type involved was unclear. Here, we showed that the VIP cell type and the way it connects to specific GC types via gap junctions are responsible for the NC between these cells.

### Activation and inhibition by VIP+ amacrine cells

We noticed that the OFF alpha-sustained GC type can be both significantly inhibited and activated by VIP+ ACs (**Fig. 1G**). This inhibition is mediated by GABA-A receptors (**Fig. 2B**), consistent with previous findings (15). One hypothesis is that this inhibition could also affect other inhibitory cells, inducing an activation effect through polysynaptic disinhibition. However, blocking GABA receptors did not fully prevent these excitatory responses (**Fig. 2B and Fig. 3B**). It is possible that, for this type, activation results from a combination of disinhibition and gap junction-mediated excitation. This behavior strikingly differed from ON alpha-sustained and ON-OFF local-edge-W3 GCs, which were only excited by VIP+ AC stimulation (**Fig. 1G**) but not inhibited, and gap junctions mediate this excitation between VIP+ ACs and GCs (**Fig. 2D and Fig. 4F**).

**Fig 2.**
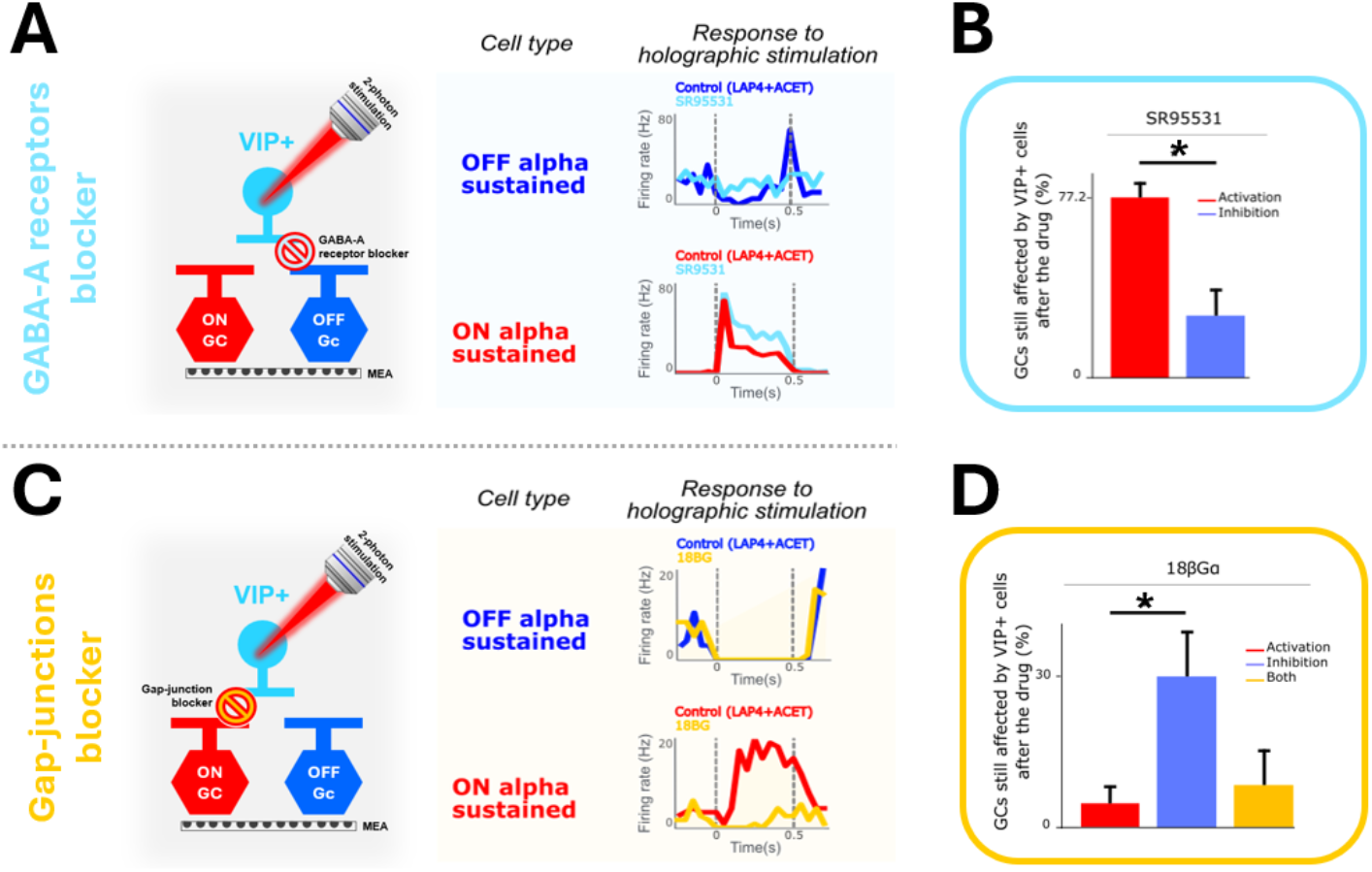
VIP+ cells excite some types of ganglion cells via gap junctions. (A, C). Effect of holographic stimulation of VIP+ cells on ON and OFF GCs after blocking GABA-A receptors (SR-95531, 20µM) or gap junctions (18BGa, 25µM). GC responses are shown in PSTHs, with the duration of VIP+ cell stimulation indicated by dashed lines (500 ms, Pmax = 0.5 mW for 1040nm). The example ON and OFF GCs are of the same type within each drug condition. (B, D). Percentage of GCs still affected by VIP+ cell stimulation after blocking GABA-A receptors and gap junctions. The values reported in the plots are mean ± S.D.M across experiments. Significance (*) is determined using a Fisher’s exact test (p < 0.001 for GABA-A blocker and p < 0.05 for 18BGa).

**Fig 3.**
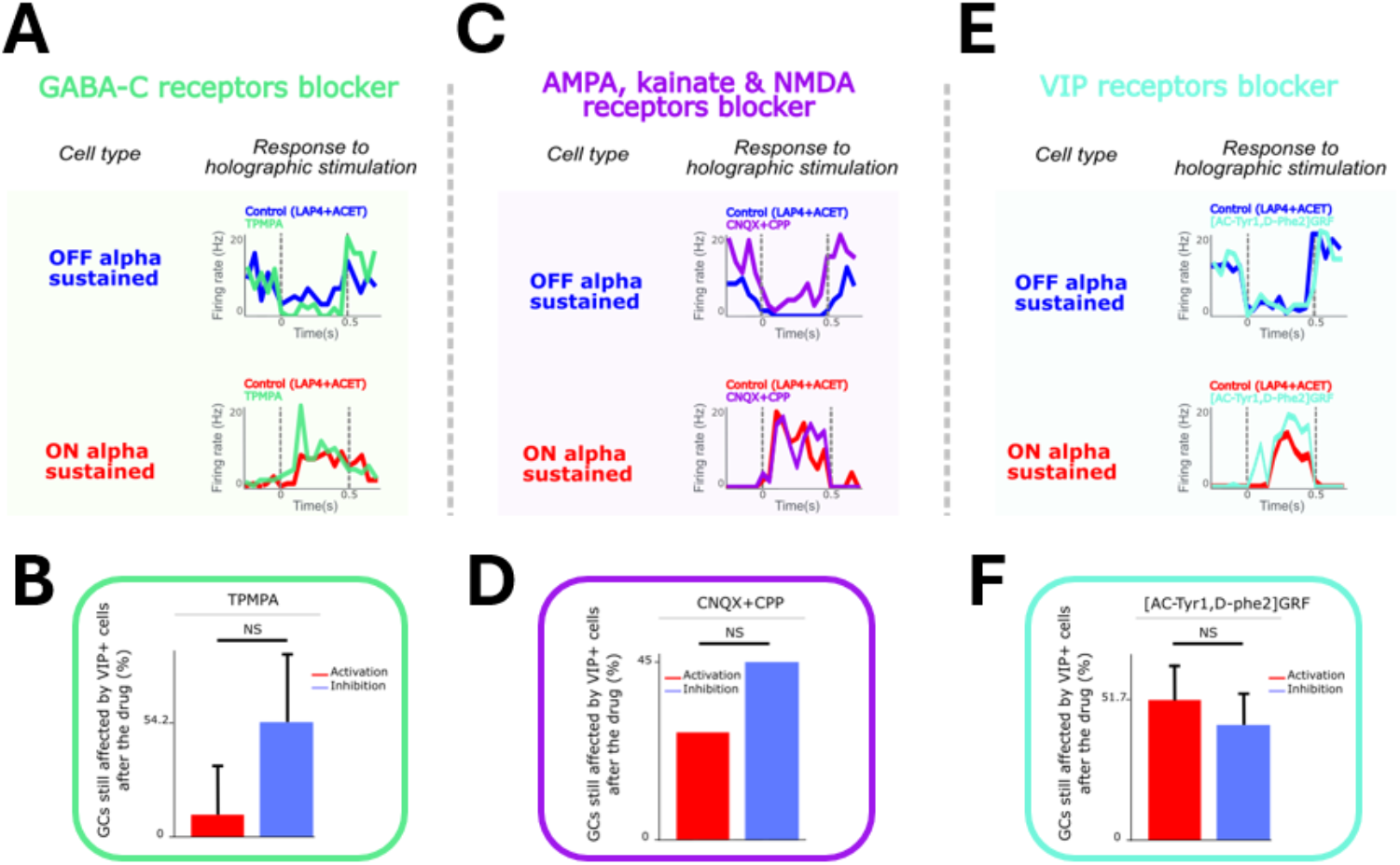
VIP+ cell excitation and inhibition effect is not mediated by other pathways than gap junctions and GABA-A receptors. (A, C, E). Effect of VIP+ cell holographic stimulation on ON and OFF GCs before and after blocking GABA-C receptors (TPMPA, 50µM), AMPA+kainate+NMDA receptors (CNQX + CPP; 20 and 10µM), and VIP receptors ([AC-Tyr1, D-Phe2]GRF, 300nM). The GCs responses before and after the application of each drug are shown in PSTHs, with the holographic stimulation duration indicated by dashed lines (500 ms, Pmax = 0.09 mW/μm2). (B, D, F). Percentages of GCs still affected by VIP+ cell stimulation after blocking GABA-C receptors, AMPA+kainate+NMDA receptors, and VIP receptors. The values reported in the plot are mean ± S.D.M across experiments. Significance was assessed using a Fisher exact test (p>0.1 for GABA-C, NMDA+AMPA, and VIP receptors).

While we focused on specific types in which we observed consistent effects, we also found that VIP+ AC stimulation could activate or inhibit other cell types. OFF step and ON-OFF-JAMB could be inhibited by VIP AC stimulation, and we also observed some activation in OFF step, ON DS transient, ON low frequency, ON-OFF JAMB, and ON-OFF DS (**Supp.Fig. 2**). Since the network of ACs is interconnected, it is also possible that some of these activations and inhibitions for these types are polysynaptic.

A limitation of our experiments is the possible lack of specificity of the pharmacological blockers we used. The gap junction antagonist used in our experiments (18BGa) might block other types of connexin in the retina or lack specificity. However, we have additional evidence that this excitation is not caused by GABAergic disinhibition, as blocking GABA-A and GABA-C receptors had no effect (**Fig. 2B and Fig. 3B**). We also showed that this excitation is independent of glutamatergic transmission (**Fig. 3D**). Our KO experiments also confirmed that gap junctions are involved in synchronizing GCs of the same type. Together, these results all converge towards our interpretation.

A noteworthy limitation of our method is that, although there are three types of VIP+ ACs in the mouse retina (15,16,22), we recorded the impact of any VIP+ cell on GCs without distinguishing them by type. Since the three types of VIP+ ACs differ in their morphology, position in the retinal layers, density across the retina, connectivity, and responses to light (14–16,22), their synaptic inputs and visual responses suggest that each VIP+ AC type may have a distinct role in visual processing in the retina. In our experiments, the injected AAV we used cannot infect a specific VIP type, and our VIP:KO:Cx36 mouse model is not selective for any particular VIP+ type (**Fig. 1B**). However, we targeted only cells in the INL, which excludes VIP-3, as they belong to the GCL. Additionally, the VIP-1 ACs are the only type that is both homologously and heterologously coupled via gap junctions (15), indicating they are most likely responsible for the GC’s excitatory responses that evoke the NC we observed (**Fig. 4D, 4E**). Since we stimulated a mix of VIP-1 and VIP-2 cells, this might induce a bit more heterogeneity in the types that are inhibited by the stimulation.

**Fig 4.**
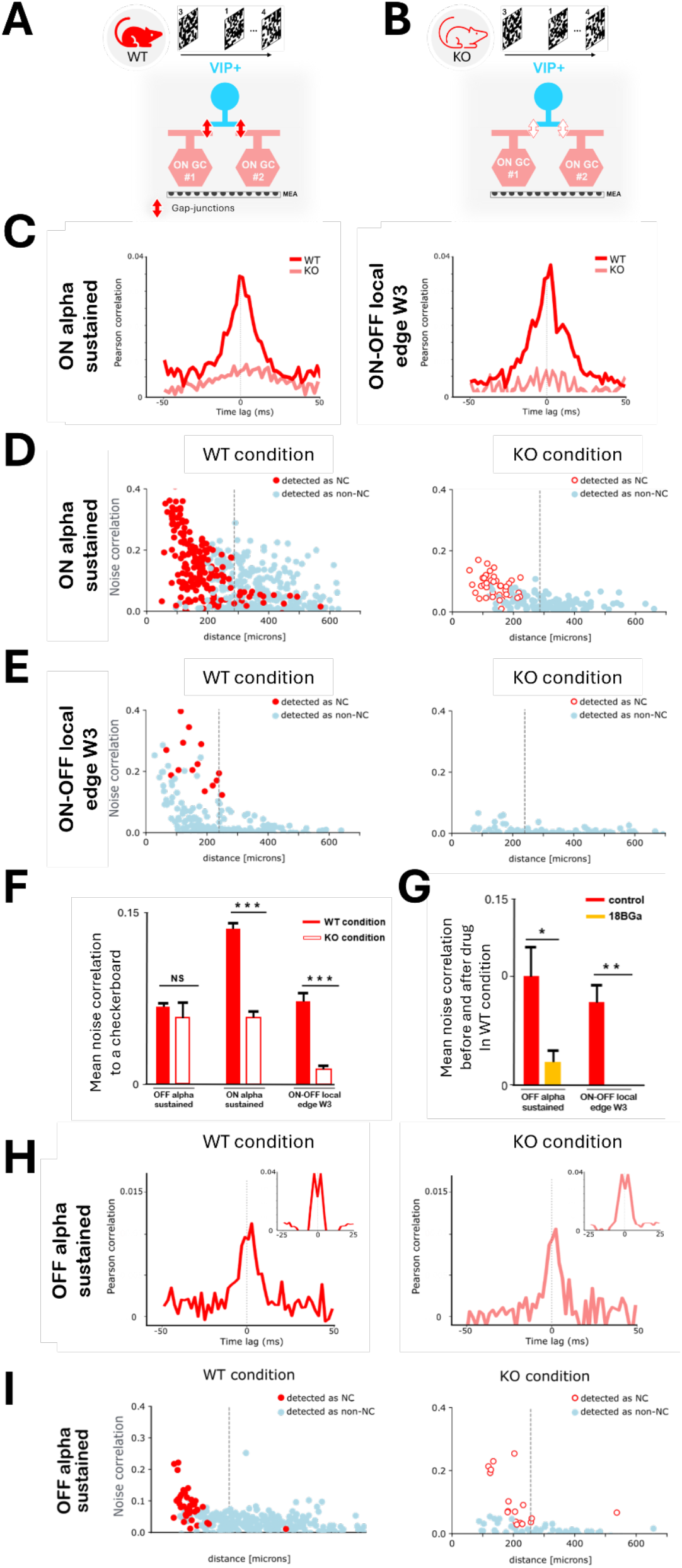
VIP+ cells can synchronize some GC types via their gap junctions. (A,B). Schematic representation of recording GCs using a multi-electrode array and visual stimulations (with a checkerboard) in WT mice (A) and VIP:KO:Cx36 mice (B). (C). Example cross-correlogram between pairs of ON-alpha-sustained GC (left) and ON-OFF local edge W3 (right) in WT (red) and KO conditions (light red). (D, E, I). Population analysis of all GC pairs of different types recorded in WT and KO mice: ON alpha sustained (n_WT_=5; n_KO_=3 retinas) (D), ON-OFF local edge W3 (n_WT_=2; n_KO_=3 retinas) (E), and OFF alpha sustained (n_WT_=5; n_KO_=3 retinas) (I). Each distribution plot shows the NC strength of each GC pair as a function of their distance (in microns) during a checkerboard. The distance between each GC pair is measured from the centers of their RFs. The grey dashed line indicates the distance within which 90% of the GC pairs with significant NC in the WT condition lie. (F, G). Comparison of the average NC intensity to a checkerboard between WT mice (F) and VIP:KO:Cx36 mice (G) for each GC type. Significance (*) is determined by using a Welch’s t-test for independent samples based on the covariance values of all GC pairs. (p < 5e-23 for ON alpha sustained, p < 5e-10 for ON-OFF local edge W3 and p > 0,1 for OFF alpha sustained.) (H). Example cross-correlogram between pairs of OFF-alpha-sustained GC in WT (left) and KO conditions (right). In the inset, the same cross-correlograms are shown with a 2 ms time bin.

### Role of the excitatory effect of VIP ACs in the retinal circuit

It has been hypothesized that the synchronized activity between GCs may be crucial for perceiving and recognizing continuous objects rather than disjointed ones (42,43). Roy et al. (2017) (43) demonstrated that a continuous object induces synchronization among widely spaced GC pairs. They show that this long-range synchronization results from gap junction connections between these GC pairs and some ACs, presumably located in between. They speculated that these ACs belong to the polyaxonal class because they are characterized by long, axon-like extensions that enable them to interact with multiple cells over long distances.

VIP-1 ACs correspond to this profile. They have multiple projections, which allow them to contact several regions of the retina (14,15,22). We demonstrated that VIP+ ACs can excite certain types of GCs (**Fig. 1G**), increase their NC, and synchronize their activity (**Fig. 4F**); they may thus play a role in global object perception, as described by Roy et al. (2017) (43). Beyond this role, this NC may also increase the information transmitted by the ganglion cell population, at least for some stimuli (44).

Our results provide new insights into VIP+ ACs in the retina, a type of AC among the more than 60 identified so far. For many AC types, their impact on GCs is unclear, making it difficult to understand their functional role. The method used in this study provides a valuable approach to understanding the role of each AC type in visual information processing as soon as genetic tools can target the cell of interest (12). It allows for dissecting the influence of individual retinal neurons on their postsynaptic circuits, revealing how they shape GC responses and contribute to retinal information processing. It also shows that the role of ACs is not restricted to lateral inhibition or gain control. VIP ACs are an example of ACs that also play a role in synchronizing nearby ganglion cells. Future functional exploration of the other AC types might reveal other new functional roles.

### Possible role of VIP+ interneurons in the cortex

VIP+ cells in the cortex appear to play a key role in regulating cortical activity based on behavioral states such as locomotion, arousal, and task engagement, as well as shaping various aspects of sensory processing and learning.

VIP+ cells influence local cortical circuits by balancing excitation and inhibition in the cortex, primarily through disinhibition and the regulation of somatostatin (SST) (45–48), which usually suppresses pyramidal neuron activity. By inhibiting SST interneurons, VIP+ cells indirectly enhance pyramidal neuron activity (41). This targeted local disinhibition helps increase the responsiveness of pyramidal neurons to relevant stimuli, thereby sharpening the tuning of cortical neurons (41,49–57).

VIP+ cells are also modulated by arousal and locomotion, and can be involved in learning (58–67). In the primary visual cortex, VIP+ cells modulate sensory processing and enhance responses to specific low-contrast visual stimuli (68). VIP+ interneurons are more active and have a stronger influence during low-contrast stimuli, such as dim or blurred objects (69), rather than high-contrast stimuli (70). VIP interneurons not only amplify all sensory inputs but also selectively boost responses to weak stimuli, particularly under low-contrast conditions, to help detect less prominent signals (69).

Given our results in the retina, it would be interesting to investigate whether VIP+ interneurons in the visual cortex also form gap junctions with other cells, whether this reshapes population activity, and whether it helps detect specific stimuli. More broadly, our results suggest that the functional repertoire of inhibitory interneurons may be more diverse than expected. First, even a single type can have a dual role. Second, and more importantly, this role cannot be fully appreciated without looking at the population level, e.g., the level of synchrony.

## Material & Methods

### Animals

All experiments were conducted on male and female mice aged 3-8 months. We used three different mouse lines. (1) A C57BL/6J mouse line from Janvier Laboratories (Le Genest-Saint-Isle, France) for wild-type control experiments. (2) A VIP-ires-Cre mouse line (VIPtm1(cre)Zjh/J; The Jackson Laboratory), in which the expression of Cre recombinase is driven by endogenous VIP regulatory elements(13). These mice were injected with adeno-associated virus (AAV) vectors to make VIP+ cells sensitive to light. This line was used during the optogenetic experiments (**Fig.1A**). (3) A VIP:KO:Cx36 mouse line, F2 offspring of VIP-ires-Cre x Cx36-fl/fl mice, specifically knockout (KO) for gap junctions in VIP+ cells. This line was used in experiments exploring the functional role of VIP+ cells in synchronization (**Fig.4A, 4B**). All animals were housed in enriched cages with free access to food and water. The ambient temperature ranged from 22 to 25 °C, the humidity ranged from 50% to 70%, and the light cycle consisted of 12–14 hours of light and 10–12 hours of darkness. All animal procedures followed the institutional Animal Care standards of Sorbonne Université. Procedures were approved by the Animal Studies Committee of Washington University School of Medicine and complied with the National Institutes of Health Guide for the Care and Use of Laboratory Animals.

### Viral injection

Viral injections were performed in the VIP-ires-Cre mouse line (heterozygous or homozygous for cre) to make them sensitive to light by expressing the optogenetic protein ReaChR for optogenetic experiments.

#### AAV production

Recombinant AAVs were produced by the plasmid co-transfection method, and the resulting lysates were purified using iodixanol gradient ultracentrifugation, as described in Dalkara et al., 2013 (71). The sequence employed is pAAV-ss-hSyn-FLEX-ReaChR-Citrine, with ReaChR linked to the green fluorescent reporter Citrine. All plasmids were produced and amplified by Vector Builder. Vector stocks were quantified by real-time PCR relative to a standard for DNase-resistant vector genomes.

#### Intravitreal injections

Mice were anesthetized with isoflurane (5% induction, 2% during the procedure). The cornea was locally anesthetized with oxybuprocaïne, and an ultrafine 30-gauge disposable needle was passed through the sclera, beneath the ora serrata, into the vitreous cavity to create a small hole and establish an entry point in the eye. A total of 1 µl of virus stock (at a concentration of 10^14 particles of AAV/ml) was injected into the vitreous body using a microsyringe (Hamilton 074487, Dutscher). The eye was moistened with Lubrithal, and Fradexam was applied afterward to promote cicatrization. Six weeks later, AAV infection was confirmed by observing fluorescent protein expression. Mice were anesthetized again with isoflurane, and mydriasis was induced with Neosynephrine and Mydriaticum to examine their eye fundus. Visualization was performed using a Micron IV device, first under white light to assess retinal structure and health, then with blue illumination and a green filter to evaluate fluorescent protein expression, thereby assessing viral infection quality and consistency.

### Electrophysiological recordings

#### MEA recording

Electrophysiological data were recorded from the isolated retinas of 25 mice in total (4 C57BL6J, 18 VIP-ire-Cre, 3 VIP:KO:Cx36). All animals were euthanized with 4% CO2 inhalation followed by cervical dislocation under dim red illumination. The eyes were enucleated and transferred into an oxygenated Ringer’s medium (NaCl, S7653, 125 mmol/L; KCl, P9541, 2.5 mmol/L; MgCl2-6H2O, M9272-5006, 1 mmol/L; NaHCO3, SSD11, 26 mmol/L; CaCl2, C3306, 1 mmol/L; L-glutamine, 53126, 0.43 mmol/L; NaH2PO4, S5761, 1.25 mmol/L; D-(+)-Glucose, 68270, 20 mmol/L, all from Sigma-Aldrich). Dissection was performed under dim light to isolate the retina from the eye cup and remove the vitreous, as described by Marre et al. (2012)(20) and Yger et al. (2018)(72). Ventral retinal pieces were used whenever possible across experiments to ensure consistency. The retina was then mounted onto a dialysis membrane coated with poly-L-lysine and lowered onto a 252-channel multi-electrode array (MEA), with the GC’s side facing first. During dissection and recordings, the tissue was constantly perfused with oxygenated Ringer’s solution (5% CO2 and 95% O2), heated to 36°C, at a rate of 3 mL/min, using a peristaltic perfusion system with two independent pumps: PPS2 (Multichannel Systems GmbH). Throughout the experiments, the room was kept in total darkness; the only light source was the stimulation itself. We used an MEA with 8 µm radius microelectrodes, spaced 30 µm apart (256MEA30/8iR-ITO, Multichannel Systems). Raw voltage traces were digitized and stored for offline analysis using a 252-channel preamplifier (MultiChannel Systems, Germany) at a sampling rate of 20 kHz.

#### Spike sorting

Spikes were isolated using SpyKING CIRCUS 1.0.628 (see Yger et al., 2018(72)). Subsequent data analysis was performed using custom Python code. We extracted the activity of a total of 2,959 neurons. We only kept cells with refractory period violations <0.5%. These constraints ensured the good quality of the reconstructed spike trains.

### Visual stimulation

Visual stimuli were presented using a white LED (MCWHLP1, Thorlabs Inc.) as the light source and a Digital Light Processing (DLP) device (DLP9500, Texas Instruments), with the light focused on the photoreceptors using standard optics and an inverted microscope (Nikon). The light level matched photopic vision: 4.9×10^4 and 1.4×10^5 isomerizations / (photoreceptor.s) for S and M cones, respectively. This luminance only activated the photoreceptors without stimulating the opsin expressed by VIP+ cells in the VIP-ires-Cre mouse line.

In all the experiments, we began by displaying three visual stimuli to identify the spatial and visual properties of each GC recorded on the MEA and to classify them by cell type. These stimuli include a checkerboard, a chirp, and a drifting grating (DG). (1) The checkerboard stimulus was used to map the receptive fields (RFs) of GCs. It consists of a white noise stimulus displayed at 30Hz for 45 minutes to 1 hour. The check size was 40 μm for optogenetic experiments and 70 μm for synchronization experiments (**Fig.1A**). To estimate the RF centers of the GCs, we computed the spike-triggered averages (STAs) of their responses to the checkerboard stimulus over 700 ms (21 frames before each spike). For each cell, this produces a 3D matrix (2 spatial dimensions and 1 temporal dimension) that represents the average check luminance across all frames preceding a spike. We extracted the spatial component of the STA by calculating its time-standardized mean across time. This yielded a 2D heat map showing which checks evoke responses in RGCs. We then fitted a 2D Gaussian distribution to this spatial component and modeled the RF center as the ellipse bounded by one standard deviation of this distribution. Cells that did not produce a good Gaussian fit were excluded from the dataset. To obtain the temporal component of the RFs, we averaged the STA across all checks within the estimated RF center. (2) The chirp stimulus was used to characterize the cell types of GCs, similar to the approach used to identify and classify 32 different mouse RGC types(23). This full-field stimulus oscillated its luminance from dark to bright (0 to 1) with a constant contrast, starting with a frequency increase, followed by a contrast increase at a constant frequency. It was presented at 50 Hz and included 20 repetitions of 32 seconds each. (3) The DG stimulus was shown at 50 Hz, moving in 8 different directions at a speed of 479.5 μm/s, with a spatial period of 959 μm (274 pixels at 3.5 μm/pix) and 50% Michelson contrast (luminance range of 0.75–0.25). Each 10-second grating was preceded by 2 seconds of gray (0.5 luminance), with a 2-second cycle period. This setup ensures each ganglion cell’s RF passes through the grating edges five times. The eight directions were repeated four times in a pseudo-random order. The stimulus profile and dynamics follow those described in Yao et al. (2018) (73) for identifying direction-selective cells, but in our case, a single luminance value was used.

### Holographic stimulation

Holographic simulations were used to optogenetically stimulate the VIP+ cells expressing the opsin ReaChR in the injected VIP-ires-Cre mice. We chose ventral retina sections because they contain the highest density of VIP type-1 and 2 cells (located in the INL) with the lowest density of VIP type-3 cells (located in the GCL), allowing us to focus the optogenetic stimulation mainly on these two VIP cell types(15,16,22). We first used one-photon fluorescence imaging to scan the AC layer and identify VIP+ cells expressing the opsin combined with a fluorescent protein. For each experiment, we selected a subset of VIP+ cells (between 10 and 25) and recorded their locations to deliver the optogenetic stimulus with high accuracy. Injected VIP+ cells showed excellent fluorescence, making it easy to distinguish their soma and trace their ramifications, which in some cases were extremely long (**Fig. 1B**).

For optogenetic stimulation, we used 2-photon digital holography(18). This technology shapes the light from an infrared laser to perform patterned 2-photon stimulation of specific cells at individual-cell resolution. Two-photon optogenetic stimulation was conducted after blocking synaptic transmission between photoreceptors and the rest of the retina, ensuring only opsin activation (see below in Pharmacology). We stimulated the selected VIP+ cells 20 times, one at a time, in a random order. Each laser stimulation lasted 500 ms at Pmax = 0.09 mW/μm^2^, followed by a 1-second pause. We used computer-generated digital holography, as detailed in Papagiakoumou et al. (2010) (74). A femtosecond pulsed beam (InSight DeepSee, Spectra-Physics) was expanded and directed onto a Spatial Light Modulator (SLM, LSH0700963, Hamamatsu). The wavelength was 1040 nm, close to the optimal range for 2p stimulation of the optogenetic protein ReaChR (975–1030 nm) (75). For this wavelength, Pmax = 0.5 mW. The SLM plane was projected onto the back focal plane of the objective lens and phase-modulated using custom software (Wavefront-Designer) to generate a precise intensity pattern at the sample plane, targeting individual opsin-expressing retinal cells. Focusing the laser on the retina, near the electrodes, causes photoelectric artifacts. This problem was addressed by high-pass filtering the recordings and then ignoring signals within a time window around the stimulation (10 ms in our case).

### Pharmacology

For optogenetic experiments that required blocking synaptic transmission between photoreceptors and the rest of the retina to stimulate only the opsin expressed by VIP+ cells, we used a combination of L-AP4 (5 µM) and ACET (1 µM), which was directly added to the perfusion medium. For experiments aimed at gaining insights into the synaptic circuitry of VIP+ cells on RGCs, we optogenetically stimulated VIP+ cells in the presence of different receptor blockers. To identify inhibitory chemical synapses, we used antagonists of GABA-A receptors (SR-95531, 20 µM) (**Fig.2A, 2B**) and GABA-C receptors (TPMPA, 100 µM) (**Fig.3A, 3B**). To block excitatory chemical synapses, we used a combination of antagonists for AMPA and kainate receptors (CNQX, 20 µM) and for NMDA receptors ([RS]-CPP, 10 µM) (**Fig.3C, 3D**). To block gap junction electrical synapses, especially those involving connexin 36, we used 18β-Glycyrrhetinic acid (18β-GA, 25 µM), which acts within approximately 20 minutes (76) (**Fig.2C, 2D**). To block VIP receptors, we used [AC-Tyr1, D-Phe2]GRF (300 nM) (29,77) (**Fig.3E, 3F**). All drugs were obtained from Tocris Bioscience, except for 18β-GA, which was from Bertin Bioreagent. All drugs were directly added to the medium to ensure delivery to the retina via perfusion. To evaluate each drug’s effectiveness, we stimulated the retina every 10 minutes from the time of drug addition to t+30 with white and dark full-field flashes of 500 ms duration at 1 Hz for 3 minutes to check the visual or optogenetic responses of GCs.

### Data analysis

#### Activation & Inhibition detection

To determine whether the GC responses evoked by holography stimulation of VIP+ cells were significant, we defined an activation and inhibition detection criterion. First, for every GC recorded, we calculated a PSTH across repetitions of holographic stimulation of VIP+ cells. We discarded GCs whose PSTH maxima were below 12Hz. Then, for the remaining PSTHs, we calculated the mean baseline activity before the holography onset (µ*b*), the mean activity and its maximum during the holographic stimulation (µ_*d*_ and *m*_*d*_), the mean activity after the holographic stimulation (µ_#_), and their respective standard deviations calculated over time and across repeats (*σb,σd,σa*)(). We first checked whether the GC had significant spontaneous activity before the holography onset (µ_*b*_ > 2*σ*_*b*_). If µ_*b*_ > 2*σ*_*b*_, we checked whether the activity during the holographic stimulation was higher than baseline: (µ_*d*_ − µ_*b*_) > 2.5*σ*_*b*_. In that case, the GC was considered activated by the holographic stimulation of the VIP+ AC. In the cases where there was no significant baseline activity before the holographic stimulation (µ_*b*_ < 2*σ*_*b*_) we checked whether the holographic stimulation activated the cell by looking at the peak of activity of the cell during the holographic stimulation (*m*_*d*_ > 2.5*σ*_*d*_). If so, we considered the cell activated by the holographic spot. For the inhibition detection, the same test was applied, except that the criterion was to be below the baseline: (µ_*d*_ − µ_*b*_) < 3.5*σ*_*b*_.

#### Pharmacology analysis

We detected GCs activation and inhibition evoked by VIP+ cells holography stimulation (see above) before and after the addition of different pharmacological drugs. For each drug, we calculated the percentage of GCs that remained affected by VIP+ cells after drug application. We used the Fisher exact test to determine whether there was a significant difference between the measured percentages before and after drug administration, as well as between the percentages of cells activated and inhibited following drug administration. The control condition included 18 retinas (2,959 good cells single units). The different pharmacological conditions included 4 retinas for gap-junction blockers, 6 for GABA-A receptor blockers, 2 for GABA-B receptor blockers, 1 for NMDA and AMPA receptor blockers, and 2 for VIP receptor blockers.

#### Noise correlation detection for WT and KO mice

We measured the noise correlation (NC) intensity and the noise covariance of GC pairs of the same type in two different retinal models: WT retinas (n=5) and KO retinas (in which the VIP+ cells were depleted of their gap junctions, n=3). Both NC and covariance were computed from GC responses to repeated sequences of a white-noise checkerboard. The GC types were determined from GC responses to the chirp stimulus and their mosaics.

We measure the noise covariance of two responses *v*_*i*_,*v*_*j*_ as:

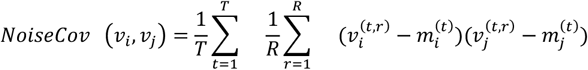

where 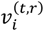 is the response of neuron i at time t and repetition r, R is the number of repeats, T is the number of time bins in the responses to a sequence, and 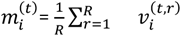.

We measured the Noise Correlations (**Fig.4F,4G**) as:

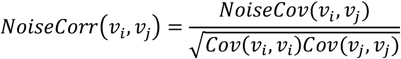

With these metrics, we built for every pair cross-covariance and cross-correlation curves by shifting one of the two responses by τ(examples of cross-correlation curves in **Fig. 4C, 4H**):

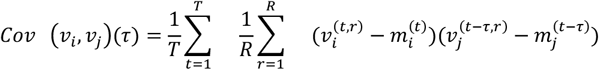

To estimate the cross-correlations and cross-covariance curves, the GC responses were binned at 2 ms.

We determined whether GC pairs were significantly correlated by examining both their cross-covariance curves and their mean firing rates. The mean rate of a cell pair was calculated as:

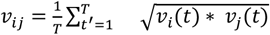

where

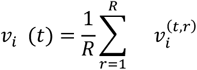

Only pairs with *v*_*ij*_> 0.06 were considered. The cross-covariance curves of the remaining pairs were smoothed with a Gaussian filter (*σ* = 3 bins ∼ 6 ms) and standardized within a time lag |τ| < 40 ms. If the peak of the standardized cross-covariance was larger than 6 S.D.M, we defined the GC pair as having a significant NC (red dots in **Fig. 4D, 4E, 4I**). Both of these controls are necessary to remove pairs that exhibit high correlation values solely due to low and noisy firing rates.

For the selected pairs, we considered the noise correlation strength of the responses, binned at 25 ms, to mitigate the effect of response noise. Finally, for each GC type, we tested whether there was a significant difference across pairs between WT and KO retinas using a Welch t-test. The difference was considered significant at p <0.05.

## Supplementary Figures

**supp Figure 1.**
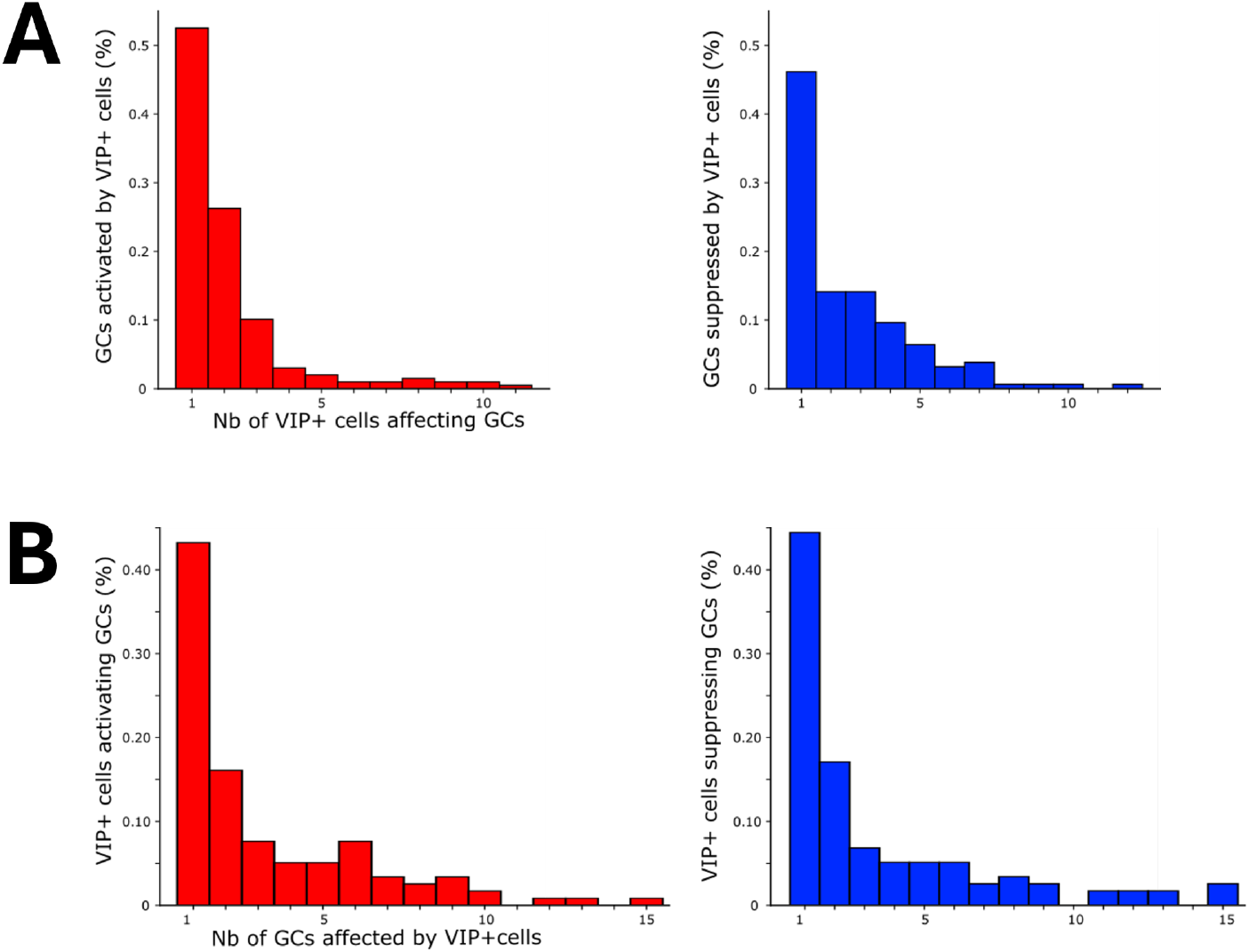
(A) Percentage of GCs affected by the holographic stimulation of different VIP+ cells. (B) Percentage of VIP+ cells that influence the activity of different GCs during holographic stimulation.

**supp Figure 2.**
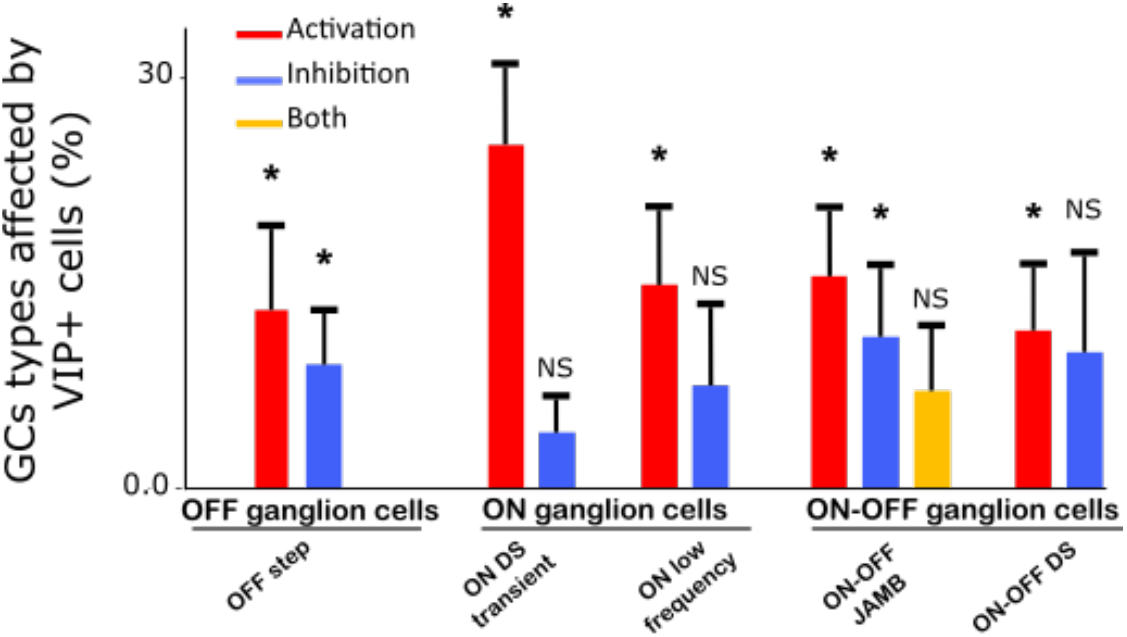
Percentages of GC types affected by VIP+ cell stimulation. The values reported in the bar plots are mean ± S.D.M. across experiments. Values are considered significant (*) if the mean is at least two *σ*s larger than zero.

## Acknowledgments

We would like to thank Victor Calbiague for help with experiments, Guihlem Glaziou for help with data analysis, Melissa Desrosiers and Deniz Dalkara for help with the vector design, and Guy Bouvier for critical reading,

## Funding

ERC Consolidator grant DEEPRETINA (101045253) (OM)

ANR grant Chaire Industrielle MyopiaMaster ANR-22-CHIN-0006 (OM)

ANR grant ANR-18-CE37-0011 –DECORE (OM)

ANR grant ANR-20-CE37-0018-04-Shooting Star (OM)

ANR grant ANR-22-CE37-0033 NUTRIACT (OM)

ANR grant project ANR-22-CE37-0016-01 PerBaCo (OM)

ANR grant project RetNet4EC (OM)

BPI funded program PREMYOM (OM)

Programme Investissements d’Avenir IHUFOReSIGHT 497 (ANR-18-IAHU-01) (OM)

Olivier Marre’s lab is part of the DIM C-BRAINS, funded by the Conseil Régional d’Ile-de-France.

## Bibliography

1. Wood KC, Blackwell JM, Geffen MN. Cortical inhibitory interneurons control sensory processing. Curr Opin Neurobiol. 2017 Oct;46:200–7.

2. Ozeki H, Finn IM, Schaffer ES, Miller KD, Ferster D. Inhibitory Stabilization of the Cortical Network Underlies Visual Surround Suppression. Neuron. 2009 May 28;62(4):578–92.

3. Adesnik H, Bruns W, Taniguchi H, Huang ZJ, Scanziani M. A Neural Circuit for Spatial Summation in Visual Cortex. Nature. 2012 Oct 11;490(7419):226–31.

4. Natan RG, Briguglio JJ, Mwilambwe-Tshilobo L, Jones SI, Aizenberg M, Goldberg EM, et al. Complementary control of sensory adaptation by two types of cortical interneurons. King AJ, editor. eLife. 2015 Oct 13;4:e09868.

5. Liu B hua, Li Y tang, Ma W pei, Pan C jie, Zhang LI, Tao HW. Broad inhibition sharpens orientation selectivity by expanding input dynamic range in mouse simple cells. Neuron. 2011 Aug 11;71(3):542–54.

6. Werblin FS. Six different roles for crossover inhibition in the retina: Correcting the nonlinearities of synaptic transmission. Vis Neurosci. 2010 Mar;27(1–2):1–8.

7. Jadzinsky PD, Baccus SA. Transformation of Visual Signals by Inhibitory Interneurons in Retinal Circuits. Annu Rev Neurosci. 2013 July 8;36(1):403–28.

8. MacNeil MA, Masland RH. Extreme Diversity among Amacrine Cells: Implications for Function. Neuron. 1998 May 1;20(5):971–82.

9. Wässle H. Parallel processing in the mammalian retina. Nat Rev Neurosci. 2004 Oct;5(10):747–57.

10. Helmstaedter M, Briggman KL, Turaga SC, Jain V, Seung HS, Denk W. Connectomic reconstruction of the inner plexiform layer in the mouse retina. Nature. 2013 Aug 8;500(7461):168–74.

11. Yan W, Peng YR, Van Zyl T, Regev A, Shekhar K, Juric D, et al. Cell Atlas of the Human Fovea and Peripheral Retina [Internet]. Neuroscience; 2020 [cited 2025 Dec 6]. Available from: http://biorxiv.org/lookup/doi/10.1101/2020.02.11.943779

12. Calbiague-Garcia V, Varro D, Buffet T, Marre O. The mysterious middlemen making your vision pop: understanding the function of amacrine cells. J Physiol. 2025;603(21):6473–502.

13. Taniguchi H, He M, Wu P, Kim S, Paik R, Sugino K, et al. A Resource of Cre Driver Lines for Genetic Targeting of GABAergic Neurons in Cerebral Cortex. Neuron. 2011 Sept 22;71(6):995–1013.

14. Zhu Y, Xu J, Hauswirth WW, DeVries SH. Genetically Targeted Binary Labeling of Retinal Neurons. J Neurosci. 2014 June 4;34(23):7845–61.

15. Park SJH, Borghuis BG, Rahmani P, Zeng Q, Kim IJ, Demb JB. Function and Circuitry of VIP+ Interneurons in the Mouse Retina. J Neurosci. 2015 July 29;35(30):10685–700.

16. Akrouh A, Kerschensteiner D. Morphology and function of three VIP-expressing amacrine cell types in the mouse retina. J Neurophysiol. 2015 Oct;114(4):2431–8.

17. Pérez-Fernández V, Milosavljevic N, Allen AE, Vessey KA, Jobling AI, Fletcher EL, et al. Rod Photoreceptor Activation Alone Defines the Release of Dopamine in the Retina. Curr Biol. 2019 Mar 4;29(5):763-774.e5.

18. Spampinato GLB, Ronzitti E, Zampini V, Ferrari U, Trapani F, Khabou H, et al. All-optical inter-layers functional connectivity investigation in the mouse retina. Cell Rep Methods [Internet]. 2022 Aug 22 [cited 2025 Dec 6];2(8). Available from: https://www.cell.com/cell-reports-methods/abstract/S2667-2375(22)00145-X

19. Spampinato G, Trapani T, Calbiague-Garcia V, Buffet T, Orendroff E, Sermet BS, et al. The Rod Bipolar Cell Pathway Contributes To Surround Responses In OFF Retinal Ganglion Cells [Internet]. bioRxiv; 2025 [cited 2025 Dec 18]. p. 2025.10.01.679777. Available from: https://www.biorxiv.org/content/10.1101/2025.10.01.679777v1

20. Marre O, Amodei D, Deshmukh N, Sadeghi K, Soo F, Holy TE, et al. Mapping a Complete Neural Population in the Retina. J Neurosci. 2012 Oct 24;32(43):14859–73.

21. Yger P, Spampinato GL, Esposito E, Lefebvre B, Deny S, Gardella C, et al. A spike sorting toolbox for up to thousands of electrodes validated with ground truth recordings in vitro and in vivo. eLife. 7:e34518.

22. Pérez De Sevilla Müller L, Solomon A, Sheets K, Hapukino H, Rodriguez AR, Brecha NC. Multiple cell types form the VIP amacrine cell population. J Comp Neurol. 2019 Jan;527(1):133–58.

23. Baden T, Berens P, Franke K, Román Rosón M, Bethge M, Euler T. The functional diversity of retinal ganglion cells in the mouse. Nature. 2016 Jan 21;529(7586):345–50.

24. Lee EJ, Park SH, Kim IB, Kang WS, Oh SJ, Chun MH. Light- and electron-microscopic analysis of vasoactive intestinal polypeptide-immunoreactive amacrine cells in the guinea pig retina. J Comp Neurol. 2002 Apr 15;445(4):325–35.

25. Bleckert A, Zhang C, Turner MH, Koren D, Berson DM, Park SJH, et al. GABA release selectively regulates synapse development at distinct inputs on direction-selective retinal ganglion cells. Proc Natl Acad Sci [Internet]. 2018 Dec 18 [cited 2025 Dec 6];115(51). Available from: https://pnas.org/doi/full/10.1073/pnas.1803490115

26. Jensen RJ. Effects of vasoactive intestinal peptide on ganglion cells in the rabbit retina. Vis Neurosci. 1993;10(1):181–9.

27. Veruki ML, Yeh HH. Vasoactive intestinal polypeptide modulates GABAA receptor function in bipolar cells and ganglion cells of the rat retina. J Neurophysiol. 1992 Apr 1;67(4):791–7.

28. Veruki ML, Yeh HH. Vasoactive intestinal polypeptide modulates GABAA receptor function through activation of cyclic AMP. Vis Neurosci. 1994;11(5):899–908.

29. Waelbroeck M, Robberecht P, Coy DH, Camus JC, De Neef P, Christophe J. Interaction of growth hormone-releasing factor (GRF) and 14 GRF analogs with vasoactive intestinal peptide (VIP) receptors of rat pancreas. Discovery of (N-Ac-Tyr1,D-Phe2)-GRF(1-29)-NH2 as a VIP antagonist. Endocrinology. 1985 June;116(6):2643–9.

30. Cunha-Reis D, Aidil-Carvalho M de F, Ribeiro JA. Endogenous inhibition of hippocampal LTD and depotentiation by vasoactive intestinal peptide VPAC1 receptors. Hippocampus. 2014 Nov;24(11):1353–63.

31. Watling KJ, Dowling JE. Effects of vasoactive intestinal peptide and other peptides on cyclic AMP accumulation in intact pieces and isolated horizontal cells of the teleost retina. J Neurochem. 1983 Nov;41(5):1205–13.

32. Zhao F, Li Q, Chen W, Zhu H, Zhou D, Reinach PS, et al. Dysfunction of VIPR2 leads to myopia in humans and mice. J Med Genet. 2022 Jan 1;59(1):88–100.

33. Bloomfield SA, Völgyi B. The diverse functional roles and regulation of neuronal gap junctions in the retina. Nat Rev Neurosci. 2009 July;10(7):495–506.

34. Völgyi B, Pan F, Paul DL, Wang JT, Huberman AD, Bloomfield SA. Gap Junctions Are Essential for Generating the Correlated Spike Activity of Neighboring Retinal Ganglion Cells. Kihara AH, editor. PLoS ONE. 2013 July 23;8(7):e69426.

35. Brivanlou IH, Warland DK, Meister M. Mechanisms of Concerted Firing among Retinal Ganglion Cells. Neuron. 1998 Mar 1;20(3):527–39.

36. VÖlgyi B, Abrams J, Paul DL, Bloomfield SA. Morphology and Tracer Coupling Pattern of Alpha Ganglion Cells in the Mouse Retina. J Comp Neurol. 2005 Nov 7;492(1):66–77.

37. Brivanlou IH, Warland DK, Meister M. Mechanisms of Concerted Firing among Retinal Ganglion Cells. Neuron. 1998 Mar;20(3):527–39.

38. Sorochynskyi O, Deny S, Marre O, Ferrari U. Predicting synchronous firing of large neural populations from sequential recordings. PLOS Comput Biol. 2021 Jan 28;17(1):e1008501.

39. Brivanlou IH, Warland DK, Meister M. Mechanisms of Concerted Firing among Retinal Ganglion Cells. Neuron. 1998 Mar 1;20(3):527–39.

40. Völgyi B, Xin D, Amarillo Y, Bloomfield SA. Morphology and physiology of the polyaxonal amacrine cells in the rabbit retina. J Comp Neurol. 2001;440(1):109–25.

41. VÖlgyi B, Abrams J, Paul DL, Bloomfield SA. Morphology and Tracer Coupling Pattern of Alpha Ganglion Cells in the Mouse Retina. J Comp Neurol. 2005 Nov 7;492(1):66–77.

42. Neuenschwander S, Singer W. Long-range synchronization of oscillatory light responses in the cat retina and lateral geniculate nucleus. Nature. 1996 Feb 22;379(6567):728–32.

43. Roy K, Kumar S, Bloomfield SA. Gap junctional coupling between retinal amacrine and ganglion cells underlies coherent activity integral to global object perception. Proc Natl Acad Sci [Internet]. 2017 Nov 28 [cited 2025 Dec 6];114(48). Available from: https://pnas.org/doi/full/10.1073/pnas.1708261114

44. Mahuas G, Buffet T, Marre O, Ferrari U, Mora T. Strong, but not Weak, Noise Correlations are Beneficial for Population Coding. PRX Life. 2025 Aug 14;3(3):033012.

45. Pi HJ, Hangya B, Kvitsiani D, Sanders JI, Huang ZJ, Kepecs A. Cortical interneurons that specialize in disinhibitory control. Nature. 2013 Nov 28;503(7477):521–4.

46. Ma S, Hangya B, Leonard CS, Wisden W, Gundlach AL. Dual-transmitter systems regulating arousal, attention, learning and memory. Neurosci Biobehav Rev. 2018 Feb;85:21–33.

47. Karnani MM, Jackson J, Ayzenshtat I, Tucciarone J, Manoocheri K, Snider WG, et al. Cooperative Subnetworks of Molecularly Similar Interneurons in Mouse Neocortex. Neuron. 2016 Apr 6;90(1):86– 100.

48. Ma S, Hangya B, Leonard CS, Wisden W, Gundlach AL. Dual-transmitter systems regulating arousal, attention, learning and memory. Neurosci Biobehav Rev. 2018 Feb;85:21–33.

49. Mesik L, Ma W pei, Li L yun, Ibrahim LA, Huang ZJ, Zhang LI, et al. Functional response properties of VIP-expressing inhibitory neurons in mouse visual and auditory cortex. Front Neural Circuits. 2015;9:22.

50. Ibrahim LA, Mesik L, Ji XY, Fang Q, Li HF, Li YT, et al. Cross-Modality Sharpening of Visual Cortical Processing through Layer-1-Mediated Inhibition and Disinhibition. Neuron. 2016 Mar 2;89(5):1031–45.

51. Khan AG, Poort J, Chadwick A, Blot A, Sahani M, Mrsic-Flogel TD, et al. Distinct learning-induced changes in stimulus selectivity and interactions of GABAergic interneuron classes in visual cortex. Nat Neurosci. 2018 June;21(6):851–9.

52. Lee AT, Cunniff MM, See JZ, Wilke SA, Luongo FJ, Ellwood IT, et al. VIP Interneurons Contribute to Avoidance Behavior by Regulating Information Flow across Hippocampal-Prefrontal Networks. Neuron. 2019 June 19;102(6):1223-1234.e4.

53. de Vries SEJ, Lecoq JA, Buice MA, Groblewski PA, Ocker GK, Oliver M, et al. A large-scale standardized physiological survey reveals functional organization of the mouse visual cortex. Nat Neurosci. 2020 Jan;23(1):138–51.

54. Keller AJ, Dipoppa M, Roth MM, Caudill MS, Ingrosso A, Miller KD, et al. A Disinhibitory Circuit for Contextual Modulation in Primary Visual Cortex. Neuron. 2020 Dec;108(6):1181-1193.e8.

55. Garrett M, Manavi S, Roll K, Ollerenshaw DR, Groblewski PA, Ponvert ND, et al. Experience shapes activity dynamics and stimulus coding of VIP inhibitory cells. Bathellier B, Gold JI, Bathellier B, Keller GB, editors. eLife. 2020 Feb 26;9:e50340.

56. Szadai Z, Pi HJ, Chevy Q, Ócsai K, Albeanu DF, Chiovini B, et al. Cortex-wide response mode of VIP-expressing inhibitory neurons by reward and punishment. Capogna M, Moore T, Ferraguti F, editors. eLife. 2022 Nov 23;11:e78815.

57. Campagnola L, Seeman SC, Chartrand T, Kim L, Hoggarth A, Gamlin C, et al. Local connectivity and synaptic dynamics in mouse and human neocortex. Science. 2022 Mar 11;375(6585):eabj5861.

58. Luo X, Guet-McCreight A, Villette V, Francavilla R, Marino B, Chamberland S, et al. Synaptic Mechanisms Underlying the Network State-Dependent Recruitment of VIP-Expressing Interneurons in the CA1 Hippocampus. Cereb Cortex N Y N 1991. 2020 May 18;30(6):3667–85.

59. Reimer J, Froudarakis E, Cadwell CR, Yatsenko D, Denfield GH, Tolias AS. Pupil Fluctuations Track Fast Switching of Cortical States during Quiet Wakefulness. Neuron. 2014 Oct 22;84(2):355–62.

60. Fu Y, Kaneko M, Tang Y, Alvarez-Buylla A, Stryker MP. A cortical disinhibitory circuit for enhancing adult plasticity. Nelson SB, editor. eLife. 2015 Jan 27;4:e05558.

61. Pakan JM, Lowe SC, Dylda E, Keemink SW, Currie SP, Coutts CA, et al. Behavioral-state modulation of inhibition is context-dependent and cell type specific in mouse visual cortex. eLife. 2016 Aug 23;5:e14985.

62. Batista-Brito R, Vinck M, Ferguson KA, Chang JT, Laubender D, Lur G, et al. Developmental Dysfunction of VIP Interneurons Impairs Cortical Circuits. Neuron. 2017 Aug 16;95(4):884-895.e9.

63. Garcia-Junco-Clemente P, Ikrar T, Tring E, Xu X, Ringach DL, Trachtenberg JT. An inhibitory pull-push circuit in frontal cortex. Nat Neurosci. 2017 Mar;20(3):389–92.

64. Kamigaki T, Dan Y. Delay activity of specific prefrontal interneuron subtypes modulates memory-guided behavior. Nat Neurosci. 2017 June;20(6):854–63.

65. Dipoppa M, Ranson A, Krumin M, Pachitariu M, Carandini M, Harris KD. Vision and Locomotion Shape the Interactions between Neuron Types in Mouse Visual Cortex. Neuron. 2018 May 2;98(3):602-615.e8.

66. Williams LE, Holtmaat A. Higher-Order Thalamocortical Inputs Gate Synaptic Long-Term Potentiation via Disinhibition. Neuron. 2019 Jan 2;101(1):91-102.e4.

67. Bastos G, Holmes JT, Ross JM, Rader AM, Gallimore CG, Wargo JA, et al. Top-down input modulates visual context processing through an interneuron-specific circuit. Cell Rep. 2023 Sept 26;42(9):113133.

68. Glickfeld LL, Histed MH, Maunsell JHR. Mouse primary visual cortex is used to detect both orientation and contrast changes. J Neurosci Off J Soc Neurosci. 2013 Dec 11;33(50):19416–22.

69. Millman DJ, Ocker GK, Caldejon S, Kato I, Larkin JD, Lee EK, et al. VIP interneurons in mouse primary visual cortex selectively enhance responses to weak but specific stimuli. eLife. 2020 Oct 27;9:e55130.

70. Cone JJ, Scantlen MD, Histed MH, Maunsell JHR. Different Inhibitory Interneuron Cell Classes Make Distinct Contributions to Visual Contrast Perception. eNeuro. 2019;6(1):ENEURO.0337-18.2019.

71. Dalkara D, Byrne LC, Klimczak RR, Visel M, Yin L, Merigan WH, et al. In vivo-directed evolution of a new adeno-associated virus for therapeutic outer retinal gene delivery from the vitreous. Sci Transl Med. 2013 June 12;5(189):189ra76.

72. Yger P, Spampinato GL, Esposito E, Lefebvre B, Deny S, Gardella C, et al. A spike sorting toolbox for up to thousands of electrodes validated with ground truth recordings in vitro and in vivo. Kleinfeld D, editor. eLife. 2018 Mar 20;7:e34518.

73. Yao X, Cafaro J, McLaughlin AJ, Postma FR, Paul DL, Awatramani G, et al. Gap Junctions Contribute to Differential Light Adaptation across Direction-Selective Retinal Ganglion Cells. Neuron. 2018 Oct;100(1):216-228.e6.

74. Papagiakoumou E, Anselmi F, Bègue A, de Sars V, Glückstad J, Isacoff EY, et al. Scanless two-photon excitation of channelrhodopsin-2. Nat Methods. 2010 Oct;7(10):848–54.

75. Chaigneau E, Ronzitti E, Gajowa MA, Soler-Llavina GJ, Tanese D, Brureau AYB, et al. Two-Photon Holographic Stimulation of ReaChR. Front Cell Neurosci. 2016;10:234.

76. Ackert JM, Farajian R, Völgyi B, Bloomfield SA. GABA blockade unmasks an OFF response in ON direction selective ganglion cells in the mammalian retina. J Physiol. 2009 Sept 15;587(18):4481–95.

77. Cunha-Reis D, Aidil-Carvalho M de F, Ribeiro JA. Endogenous inhibition of hippocampal LTD and depotentiation by vasoactive intestinal peptide VPAC1 receptors. Hippocampus. 2014 Nov;24(11):1353–63.

